# Neural correlates of changing food choices while bypassing values

**DOI:** 10.1101/2022.06.27.497771

**Authors:** Anoushiravan Zahedi, Sergio Oroz Artigas, Nora Swaboda, Corinde E. Wiers, Kai Görgen, Soyoung Q. Park

## Abstract

Current theories suggest that altering choices requires value modification. To investigate this, normal-weight participants’ food choices and values were tested before and after an approach-avoidance training (AAT), while neural activity was recorded during the choice task using functional magnetic resonance imaging (fMRI). During AAT, participants consistently approached low-while avoiding high-calorie food cues. AAT facilitated low-calorie food choices, leaving food values unchanged. Instead, we observed a shift in indifference points, indicating the decreased contribution of food values in food choices. Training-induced choice shifts were associated with increased activity in the posterior cingulate cortex (PCC). In contrast, the medial PFC activity was not changed. Additionally, PCC grey matter density predicted individual differences in training-induced functional changes, suggesting anatomic predispositions to training impact. Our findings demonstrate neural mechanisms underlying choice modulation independent of valuation-related processes, with substantial theoretical significance for decision-making frameworks and translational implications for health-related decisions resilient to value shifts.

## Introduction

Optimizing dietary patterns is essential for aiding humans^1-4^ and global environmental health^5-7^. Current theories assume that changing choices are accompanied by modifying values associated with targeted items^8-11^. However, the necessity of shifting values for altering choices has recently become a matter of intense debate^12-19^. Whether choice can be affected regardless of value has substantial theoretical significance for understanding the cognitive and neural mechanisms of decision-making^11,19^ and translational implications for strategies used to optimize health-related decisions, such as food choices in clinical and normal populations^20-23^.

Choices are simultaneously affected by multiple factors^8-11^. The most intuitive feature is the expected reward value associated with possible options, leading to choosing the item with the highest subjective value^24-27^. Alternatively, choices can also be affected in the absence of value modification, a procedure that is largely uninvestigated in terms of underlying cognitive processes and neural substrates^11^. For example, direct training to approach certain food cues could lead to a higher frequency of choosing the targeted item, independent of the value associated with the response^28^. Notably, the difference between value- and non-value-based mechanisms is not related to their automaticity. That is, once well-trained, both mechanisms can lead to automatic responses^24,29^ that are not resource-consuming^30,31^. Additionally, if enough cognitive resources are available, both mechanisms can be overridden by using inhibition^32^. Still, since inhibition, like other forms of cognitive control, requires an immediate mental effort, it is aversive. Consequently, we prefer to rely on cognitive control processes as little as feasible^33,34^, highlighting the importance of regiments that can affect choices without resorting to inhibition^21,22,35,36^.

One approach to change choices in the absence of value modification is employing approach-avoidance training (AAT)^37,38^, in which targeted stimuli are consistently associated with approach or avoidance. For instance, Wiers et al. ^37^ showed that a short AAT, associating alcoholic beverages with avoiding response, induced an avoidance bias towards alcohol in alcohol-dependent participants. Importantly, this avoidance bias was accompanied by symptom improvement in these patients. AAT can be considered a nonreinforced learning procedure^18^ since the targeted stimuli are not connected to reward or punishment contingencies but to specific responses. Consequently, even though AAT modifies behavior, it is largely unknown how this is related to value. Hence, AAT, a canonical training regime for affecting choices, is highly appropriate for assessing the necessity of shifting values when altering choice behavior.

Although the neural underpinnings of AAT in the healthy population remain uninvestigated^39,40^, the neural substrates of values and choice behavior allow us to draw hypotheses. In the last decades, the ventromedial prefrontal cortex (vmPFC), orbitofrontal cortex (OFC), and posterior cingulate cortex (PCC) have been shown to be associated with choices^41,42^. Notably, tasks that target final choices seem to strongly engage PCC^43,44^, but value signals are represented in vmPFC and OFC^45,46^. Additionally, PCC has been discussed as the neural hub that detects changes in the environment and motivates shifts in behavior^47^. For instance, Kable and Glimcher ^43^ found that PCC activity can predict participants’ final choices in an intertemporal choice task. Also, Cousijn et al. ^48^ showed that approach bias towards cannabis-related images correlates with increased activity in PCC when comparing heavy-users and non-users. Further, PCC and precuneus have been shown to be more activated in internet-gambling-dependent participants than in the control group, and their BOLD activities are correlated with their gaming urge^49^. Together, these results suggest that compared to vmPFC and OFC that encode values and integrate them in decision-making-related networks, PCC activity might encode and incorporate internally-motivated^50^ behavioral tendencies in decision-making processes.

In the current study, we investigate the necessity of shifting values for altering choices. Further, we elucidate how such a choice modification without value-change is potentially incorporated into decision-making networks at the neural level. We hypothesize that even though AAT affects choices, it leaves reward value unaffected. Further, we assess whether the relationship between choices and ratings is modulated by AAT such that the same values would be less critical for final choices after AAT compared to baseline^51,52^. Accordingly, we expect that at the neural level, mPFC that encodes values would not be affected by AAT. In contrast, PCC, which is the neural hub that incorporates behavioral tendencies in choice behavior, would be modulated by AAT. We further explore anatomical predispositions to behavioral modifications by conducting voxel-based morphometry. Finally, to test if the AAT training effect can be transferred to real-life food intake, participants underwent a breakfast-buffet test at the end of each testing session (Fig. 1).

**Figure 1.**
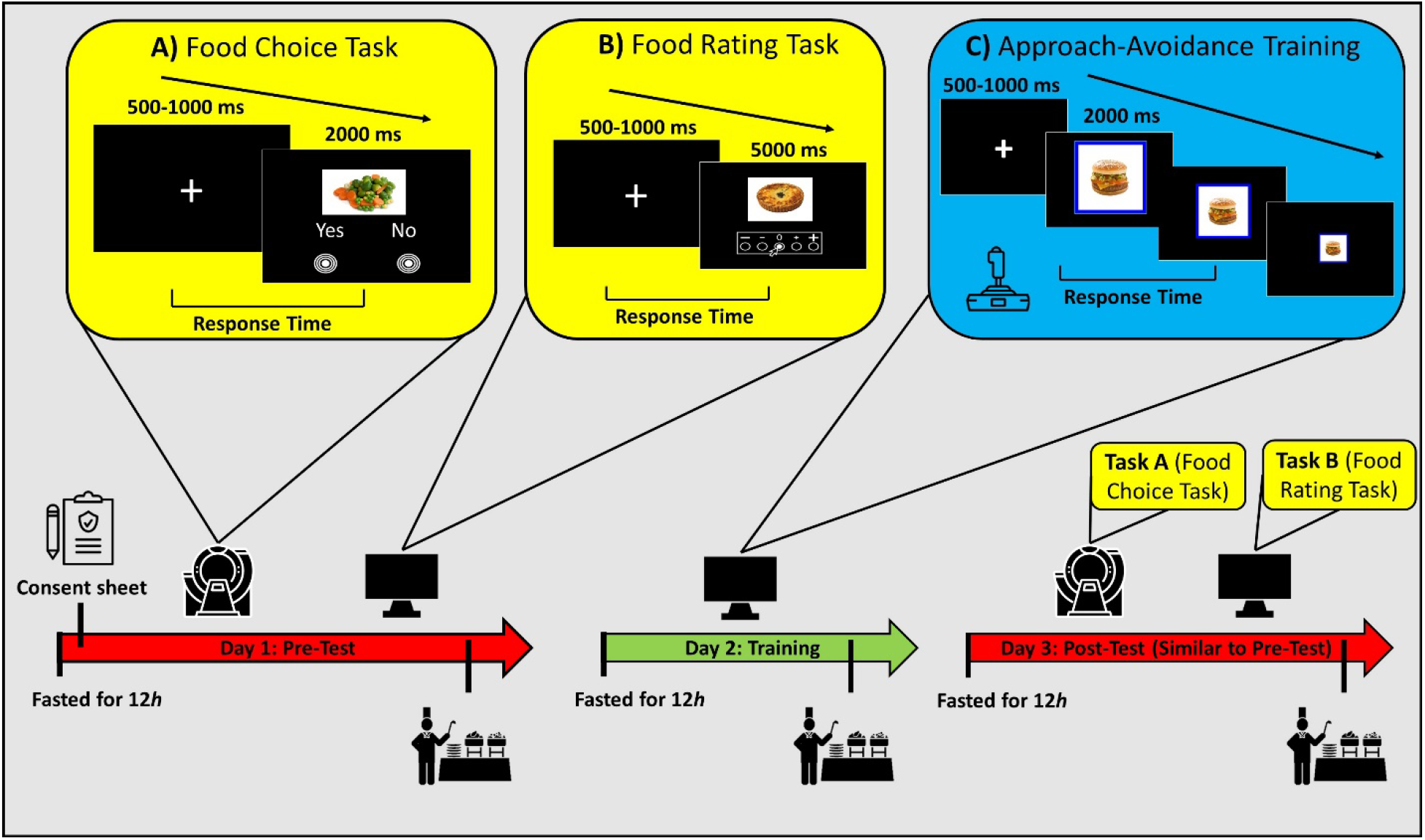
Schematic representation of the experimental paradigms. As stimuli, 80 food images^53^, containing 40 low- and 40 high-calorie food items, were employed. To test training effect generalizability, two sets of food images, each containing 20 low- and 20 high-calorie food cues, were used; one exclusively for the pre- and post-tests and the other for AAT (i.e., the training session). During Days 1 and 3, subjects participated in a food choice task, while neuroimaging data were recorded using fMRI, and a food rating task, which took place outside of the scanner. A) In the choice task, participants indicated whether they wanted to consume the presented food. The choice task consisted of 160 trials, divided into four blocks of 40 trials. The order of stimuli was pseudorandomized so that in each block, each stimulus would be presented once. B) In the rating task, participants were instructed to indicate how appealing they find the presented picture using a continuous slider. All images in the test set were rated using a continuous scale. C) For AAT, participants were instructed to consistently pull [towards their body] or push [away from their body] the presented food items in response to the color of stimulus frames (i.e., blue or yellow) using a joystick. Notably, low-calorie stimuli were consistently cued to be approached and high-calorie stimuli to be avoided. For optimal approach and avoidance resemblance^54^, the employed AAT had an embedded zooming feature. AAT consisted of five blocks of 40 trials (each picture in the training set was presented five times). The order of images was pseudorandomized so that each image would be presented once during each block. Additionally, participants were instructed not to drink or eat anything besides water for 12 hours before each session. At the end of each session, they could choose as many items as they wanted from a breakfast buffet where multiple food options were available.

## Results

We investigated 34 normal-weight female participants (*N* = 34, mean age = 25.14 years, SD = 4.01 years; mean BMI = 21.46, SD = 1.77) on three consecutive days in a within-subject design consisting of pre-training, AAT, and post-training sessions (Fig. 1). In the pre- and post-training sessions (day 1 and 3, respectively), participants performed (1) a food-choice task, where they accepted or rejected the presented food picture, while their brain activity was recorded using fMRI. (2) Subsequently, participants rated the same food items outside of the scanner. On the second day during AAT, low- and high-calorie food items were presented in colored frames (i.e., blue and yellow), and participants were instructed to avoid foods framed in one color (e.g., yellow) and approach the other (e.g., blue) using a joystick. In fact, frame colors were consistently associated with low- and high-calorie items, thereby making high-calorie items always avoided and low-calorie items approached. Participants were instructed to fast for 12 hours before each session (i.e., do not drink or eat anything besides water). After each session, they were served breakfast where they could choose as many items as they wanted from multiple food options.

### The Impact of AAT on Choices and Values

In order to investigate whether AAT affected food choices, we tested the acceptance rates via a 2 × 2 ANOVA with Calorie (low-vs. high-calorie) and Session (pre-vs. post-test) as within-subject factors. The interaction between Session and Calorie was significant (*F*(1,32) = 6.22, ***p* = 0.017** 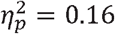, suggesting a significant change in choice as a function of training (Fig. 2-A). Notably, after the training, participants chose low-calorie food items significantly more often (*t*(32) = 3.56, ***p* = 0. 001**, *Cohen’s d* = 0.62, Bonferroni-corrected), whereas the high-calorie items were not significantly affected by AAT (*t*(32) < 1, *n. s*). Further, the main effect of Session was marginally significant (*F*(1,32) = 3.66, *p* = 0.064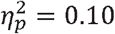. Also, participants chose significantly more low-calorie (*mean* = 0.633, *SD* = 0.190) than high-calorie (mean = 0.435, SD = 0.220) food items, as shown in significant main effect of calorie (*F*(1,32) = 12.00,***p*** < 0.001 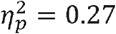.

**Figure 2.**
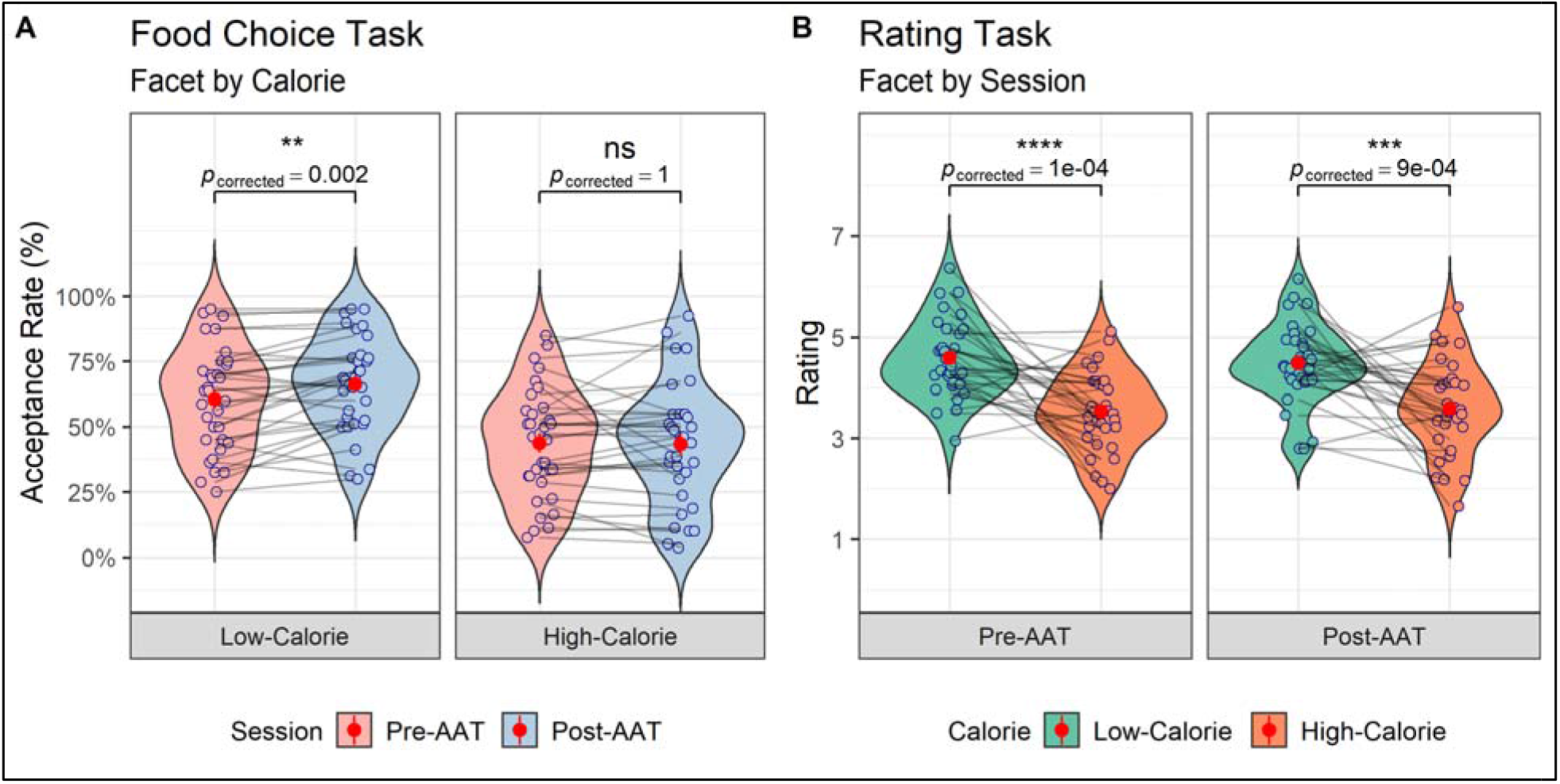
The violin diagrams of the behavioral results for (A) the food-choice task and (B) the subjective ratin task. Red dots and lines represent means and 95% confidence intervals (CI), calculated based on the standard error of the mean (SEM). All *p* values are Bonferroni-corrected.

Next, we tested whether the values associated with the food items changed after training (Fig. 2-B). Importantly, neither the interaction between Session and Calorie was significant (*F*(1,30) = 1.23, *p* = 0.27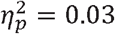, nor the main effect of Session (*F*(1,30) < 1, *n. s*.), showing that AAT did not affect food values. In general, participants rated low-calorie food items (*mean* = 4.53, *SD* = 0.78) significantly higher compared to high-calorie food items *mean* = 3.55 *SD* = 0.85; *F*(1,30) = 21.65, ***p* < 0.001**(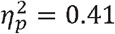.

Afterward, to assess whether AAT effects are transferred to real-life food choices, we analyzed the breakfast choices using a quasi-Poisson mixed model with Calorie (low-vs. high-calorie) and Session (pre-vs. post-test) as fixed and Participant as a random effect. We could find a marginally significant change in real food choice as a function of training, as unveiled by the marginally significant interaction of Session and Calorie (*χ*^2^(1, *N* = 132) = 3.28, *p* = 0.069). In general, participants chose more low-(*mean* = 6.72, *SD* = 2.29) than high-calorie items (*mean* = 4.69, *SD* = 2.37, χ^2^(1, *N* = 132) = 38.57, p < 0.001). We did not observe a significant main effect of Session (*χ*^2^(1, *N* = 132) = 0.31, *p* = 0.576). Notably, low-calorie breakfast choices were numerically increased (*mean* = 0.36, *SD* = 2.07), and high-calorie ones decreased (*mean* = −0.72, *SD* = 2.73) in post-compared to pre-test. However, these numerical changes, either for low-calorie (*Wilcoxon*(32) = 68, *p* = 0.168, *r* = 0.24, Bonferroni-corrected) or high-calorie breakfast choices (*Wilcoxon*(32) = 258, *p* = 0.211, *r* = 0.20, Bonferroni-corrected), were not significant.

### The Contribution of Values to Choice Shifts as a Function of Training

Next, we tested how food values contribute to choices before and after AAT. To do so, we employed individual indifference points, which indicate the estimated value at which participants chose food items with a 50% probability^51,52^. Here, we observed a significant interaction between Session and Calorie (, indicating that indifference points were modulated as a function of training. The contrast analysis (Fig. 3-A, B) showed that although there was no significant difference between low and high-calorie indifference points in the pre-training session (, Bonferroni-corrected), this difference became significant in the post-training session (, Bonferroni-corrected). Further, although high-calorie indifference points were not changed after AAT (), the low-calorie indifference points decreased significantly (, Bonferroni-corrected). These findings suggest that after training, novel behavioral tendencies were formed, which resulted in low-calorie food choices becoming less dependent on reward values.

**Figure 3.**
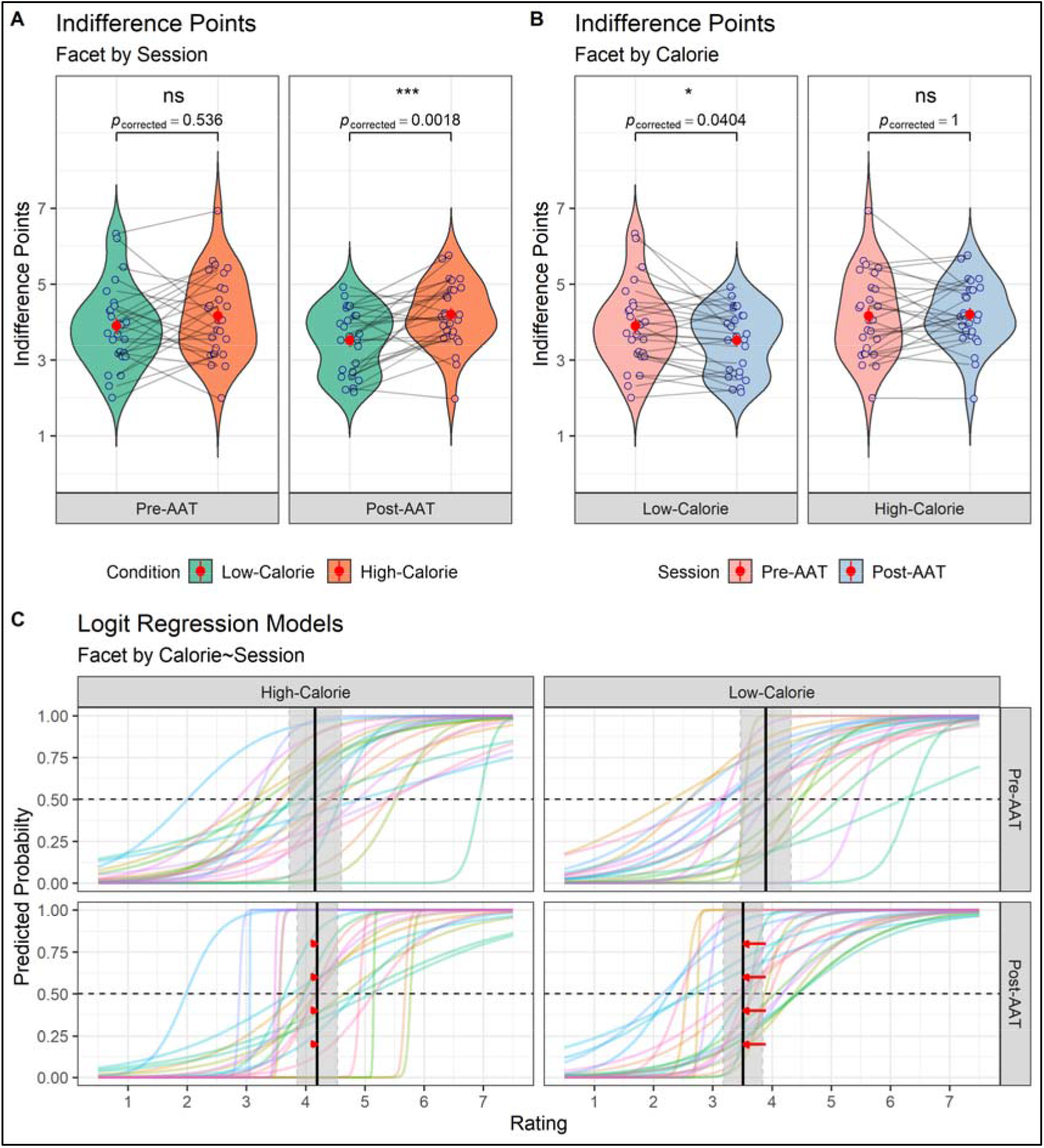
The violin diagrams of the indifference point analysis grouped by (A) Calorie and (B) Session. Red dots and lines represent means and 95% CIs, calculated based on the standard error of the mean (SEM). All *p* values are Bonferroni-corrected. (C) The predicted probabilities based on logit regression models. Dashed horizontal lines mark 50%, predicted probability, used for calculating indifference points for each participant and condition. Solid vertical lines mark means indifference points, and grey highlighted areas represent 95% CIs, calculated based on the standard error of the mean (SEM). Red arrows in the post-training panels show the changes in mean indifference points from pre to post-AAT sessions.

Further, the indifference points were not significantly changed by Session (*F*(1,25) = 1.52, *p* = 0.229,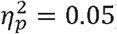. However, we observed significantly lower indifference points for low-(*mean* = 3.70, *SD* = 0.98) vs. high-calorie images (*mean* = 4.17, *SD* = 0.99) in general (*F*(1,25) = 6.45,***p* = 0.017**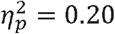. This result might indicate that participants’ choices of high-calorie compared to low-calorie food cues were influenced by other factors, such as inhibition.

To confirm that alterations in acceptance rates were related to changes in indifference points, we calculated the repeated-measure correlation coefficient between them. The results showed a strong correlation between the acceptance rates and indifference points (*r*(77) = -0.688, 95% *CI* [-0.790 -*0*.*549*, ***p****<.001*), confirming that alterations in acceptance rates were indeed associated with changes in indifference points.

### Identifying Brain Regions of Interest Involved in Value and Choice

We first identified brain regions that code the caloric value^20,55^ (i.e., high-vs. low-calorie) in th pre-test. Two areas showed significant changes: PCC [*MNI*: *x* = −3, *y* = −16, z = 35, (*t*(32) = 5.47, ***FWC***_***c***_ **=0. 035**)], and ACC [*MNI* = *x* = 9, *y* = 26, z = 26, (*t*(32) = 4.60, ***FWE***_***c***_ **=0. 007**)]. Next, we tested for decision-sensitive regions (i.e., accept versus reject) in the pre-test. In line with the previous findings^41,42,45,46^, mPFC was identified as the only region reflecting choice [*MNI* = *x* = 6, *y* = 41, z = −7, (*t*(32) = 6.29, ***FWE***_*c*_ =0. 015)]. The following analyses will thus specifically target the activities within these regions of interest.

### The Effects of Behavioral Modifications on Brain Activity

To address whether brain activity was modulated by the experimental manipulations, we analyzed the BOLD activities of the functional ROIs, which were specified in the whole-brain analysis. Using general linear models (GLM), we included Decision (Accept vs. Reject), Calorie (high-vs. low-calorie), and Session (pre-vs. post-test) as within-subject regressors.

In the mPFC ROI, the main effect of Decision was significant as expected^41,42,46^ (*F*(1,32) = 22.95, ***p*** < 0.001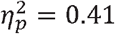, suggesting that mPFC^45,46^ is involved in choice behavior. However, mPFC activity was not significantly affected by Calorie (*F*(1,32) = 3.12,*p* = 0.086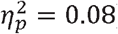, Session (*F*(1,32) = 3.16, *p* = 0.084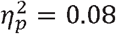, or by interactions between the factors (*Fs*(1,32) < 1, *n. s*.).

Next, we turned to PCC to investigate its role in the decision-making network. In the PCC ROI (Fig. 5), the interaction between Calorie and Session (Fig 5-A, B) was significant (*F*(1,32) = 4.94,***p* = 0.033**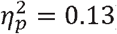, indicating that PCC activity was modulated as a function of AAT. The contrast analysis showed that although activity in response to low-compared to high-calorie food items was significantly lower in the pre-training session (*t*(32) = 5.05, ***p*** < 0.001, *Cohen’s d* = 0.87, Bonferroni-corrected), there was no significant difference between them after AAT (*t*(32) < 1, *n. s*.). Further, even though AAT significantly affected activity in response to low-calorie images (*t*(32) = 2.46, ***p* = 0. 038**, *Cohen’s d* = 0.42, Bonferroni-corrected), it did not significantly affect high-calorie items (*t*(32) < 1, *n. s*.). Together, these results suggested that PCC is a strong candidate for integrating newly formed behavioral tendencies in choice behavior, which leads to an increase in the frequency of choosing low-calorie food items. To confirm this postulation, we investigated whether the activities in response to low-calorie food items were correlated with the acceptance rate of these food images by calculating a repeated-measure correlation. Notably, the results revealed a significant correlation between PCC activity and low-calorie acceptance rates (*r*(32) = 0.352, 95% *CI* [0.003, 0.624], ***p* =. 041**; Fig. 5-C).

**Figure 4.**
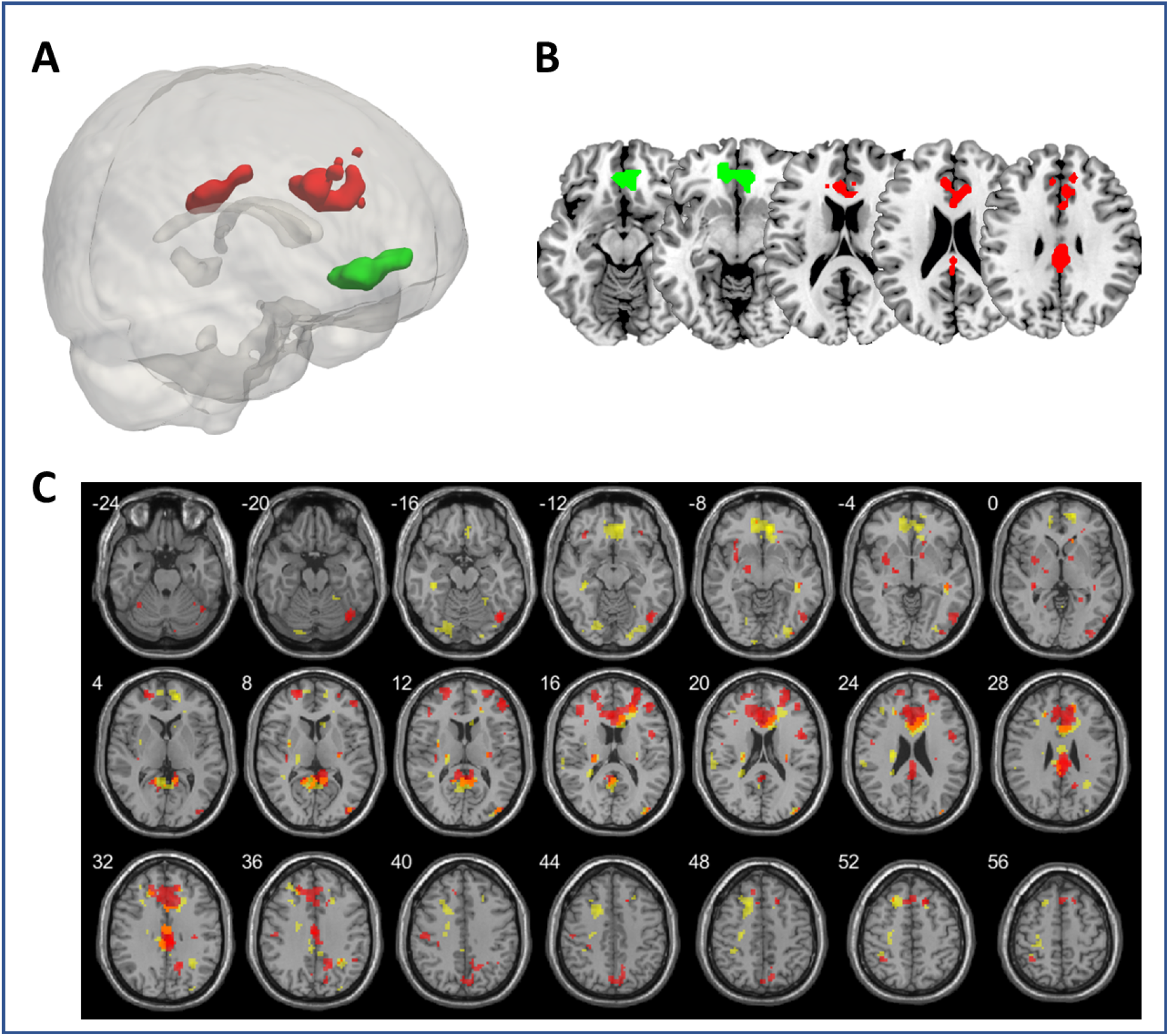
Functional ROIs derived from high vs. low-calorie (red) and accept vs. reject (green) contrasts in the pre-training session. A) The 3D rendition of the ROIs. B) The multiple-slice rendition of axial slices at z = −12, −8, 18, 22, and 30. Green: decision-sensitive area (i.e., mPFC); Red: habit-sensitive areas (i.e., ACC and PCC). C) The corresponding whole-brain analyses of high vs. low-calorie (red) and accept vs. reject (yellow). For visualization, the uncorrected images (*p* < 0.005, *k* > 10) are shown^56^.

**Figure 5.**
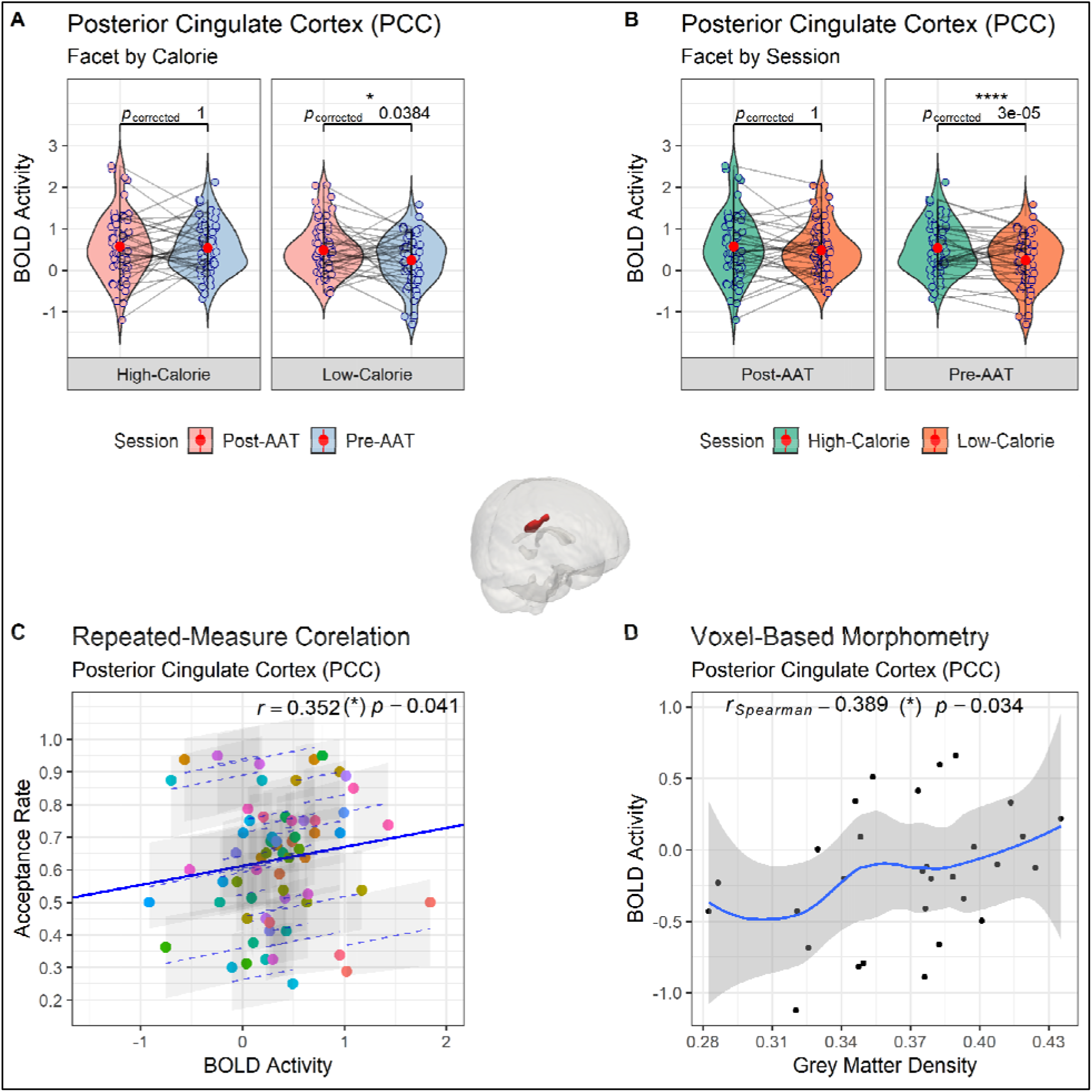
The violin diagrams of the BOLD activity of PCC faceted by (A) Calorie and (B) Session. Red dots an lines represent means and 95% CIs, calculated based on the standard error of the mean (SEM). All *p* values are Bonferroni-corrected. (C) Correlation between the BOLD activity of PCC in response to low-calorie food items and the acceptance rate of low-calorie food items in the food-choice task. (D) Correlation between grey matter densit and BOLD activity of PCC in response to low-calorie food items (depicted for 29 participants in the range:. Blue lines and grey areas show the fitted curve and 95% CI, respectively.

Additionally, Calorie significantly affected PCC activity (*F*(1,32) = 6.85,***p* = 0.013**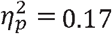, possibly showing that, in general, choosing high-calorie food items relies more on behavioral tendencies than low-calorie items^20,55^. No other main effect (Session:(*F*(1,32) = 1.81,*p* = 0.186 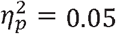 Decision: (*F*(1,32) = 3.03, *p* = 0.090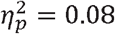 or interaction (Decision * Session: (*F*(1,32) = 2.79,*p* = .104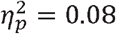 others: *Fs*(1,32) < 1, *n. s*.) was significant.

Finally, in the ACC, the interaction between Calorie and Session was marginally significant (*F*(1,32) = 3.72,*p* = 0.062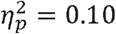, suggesting that AAT possibly affected the activity of ACC. To understand this marginal interaction, we conducted a series of contrast analyses. The results showed that even though BOLD activity in response to low compared to high-calorie food items was significantly lower before AAT (*t*(32) = 4.66, p < 0.001, *Cohen’s d* = 0.81, Bonferroni-corrected), there was no significant difference between low and high-calorie BOLD activity after AAT(*t*(32) < 1, *n. s*.). However, there was no significant repeated-measure correlation between the ACC BOLD activities and acceptance rates (*r*(32) = 0.024, 95% *CI* [−0.327, 0.369 ], *p* = .891). Further, the main effect of Calorie was significant (*F*(1,32) = 9.44,***p* = 0.004**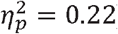. However, no other main effect (Decision: (*F*(1,32) = 2.96,*p* = 0.094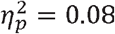 Session: *F*(1,32) < 1, *n. s*.) or interaction was significant (Session * Decision: (*F*(1,32) = 1.90,*p* = 0.176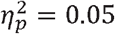, others: *Fs*(1,32) < 1, *n. s*.).

### Understanding Individual Differences in Brain Activity Alterations based on the Anatomical Predisposition

After finding that PCC might play an essential role in integrating behavioral tendencies in choice behavior, we were curious whether anatomical differences in PCC can potentially explain individual differences in brain activity changes. VBM results showed that there is a significant correlation between activity changes from the pre-to post-test and the grey matter density of PCC (*r*_*Spearman*_(31) = 0.353 [0.011, 0.621], ***p = 0. 044*)**. Notably, applying more conservative outlier criteria (]) did not change the results. (; Fig. 5-D).

## Discussion

Current decision-making theories assume that changing choices are dependent on modifying values associated with targeted items^8-11,18^. In the current study, we challenge this assumption by investigating the effects of AAT on choice behavior, during which low-calorie food items were consistently associated with approach and high-calorie food items with avoidance. The behavioral results showed that even though subjective ratings of food items were not changed after AAT, participants chose low-calorie food items more frequently. Further, logit regression modeling of indifference points revealed that the contribution of values in final choices was significantly reduced after training. Hence, our behavioral results strongly corroborated our hypothesis that AAT modulates choices without value modification. The fMRI results indicated that BOLD activity of PCC, but not mPFC, was modulated as a function of AAT. Notably, PCC activity in response to low-calorie images during pre- and post-tests was correlated with participants’ acceptance rates, indicating a fundamental role of PCC in integrating behavioral tendencies in choice behavior.

Our behavioral findings are in line with replicated findings in the AAT^12-15,38,39,57,58^ and other nonreinforced learning literature^16,59-61^. For instance, Kakoschke et al. ^14^ found that AAT decreases unhealthy food choices compared to cognitive-control training (where a go-nogo task was used to increase inhibitory control for unhealthy food cues) and the control condition. However, in their study, AAT did not affect food items’ implicit associations (IAT). Another form of nonreinforced learning is cue-approach training. In cue-approach training, targeted stimuli [e.g., low-calorie food images] are consistently associated with a cue [e.g., a high pitch tone], which signals that the presented item should be chosen [e.g., pressing a button]. Multiple studies showed cue-approach training boosts choices of trained items^16,59-61^, although the increase in choices is not accompanied by an increase in subjective ratings of the targeted items^16,17,59^.

Notably, our results showed that alterations in indifference points were correlated with choice shifts. When one considers that the indifference points of low-calorie items are the only differentiating variable between the pre- and post-AAT, our suggested account becomes even more credible. That is, our results indicate that alteration in choices is the consequence of the formation of novel behavioral tendencies rather than shifting values associated with the targeted items. This suggestion is supported by findings in the cue-approach training literature that indicate the consciously perceivable association between specific stimuli and responses is crucial for the effectiveness of nonreinforced learning^16,17^.

Secondly, our results indicated that the effects of food-related AAT in normal-weight participants are restricted to elevating the choices of low-calorie items. This finding is in line with other studies; for instance, Mehl et al. ^13^, using a multi-session training design, compared obese and normal-weight subjects. They observed that in the healthy group, AAT only increased the approach bias towards healthy food items; in contrast, in the obese group, AAT only decreased the approach bias towards unhealthy food images. Assessing the impact of other nonreinforced learnings, such as cue-approach training, offers a similar pattern. For instance, even though cue-approach training can increase the frequency of choosing liked items, it cannot decrease the frequency of selecting neutrally or positively-valenced items^16-18,61^. These results can be understood if one considers that behavioral tendencies are one variable that affects choices and interacts with other factors such as value associations. Although a short AAT session can increase choices of approach-trained items that are appetitive or positively valenced, it would not decrease choices of avoid-trained images when they are appetitive or liked. It is still feasible that multiple-session or longitudinal AAT would negate the effects of other factors, such as existing values. The distinction between values and behavioral tendencies is further corroborated by the effects of evaluative conditioning^62^, where it is repeatedly shown that changes in preferences are not necessarily accompanied by alterations in the choice behavior^63-65^.

By assessing participants’ breakfast choices, we investigated the translational effects of AAT on real-life food choices. Our results showed that participants’ breakfast choices are modulated by AAT depending on the calorie content of food items, highlighting a numerical increase in low-calorie and decrease in high-calorie choices in the post-compared to the pre-test. Considering these results, one might infer the translational effects of AAT on real-life decisions. However, the results of previous studies investigating AAT effects on real-life food choices are mixed, as some show that AAT can significantly affect real-life choices^12^, but others failed to find these effects^15,57^. Considering the mixed findings by other groups and our marginally significant results, one should cautiously interpret these results before future studies with larger samples investigate the reliability and stability of these effects.

Our fMRI results not only corroborate the behavioral modification account but also provide a new framework for understanding previous fMRI results. Considering that in our results, there is a correlation between acceptance rates of low-calorie images and the activity of PCC, one can suggest PCC as the neural hub that integrates behavioral tendencies during decision-making. This proposition is entirely in line with value-based decision-making literature, where it is shown that PCC activity is especially crucial in tasks that measure choices^41-44^ but might not be engaged in tasks that measure values and preferences such as willingness to pay^45,46^. Especially, the observed results highlight the significance of PCC activity in behavioral modifications, as a similar neural pattern can be observed in other nonreinforced learning procedures. The increase in the frequency of choosing an item is correlated with the increase in the activation of posterior parietal areas, including PCC and precuneus^48,66,67^, and frontal areas such as ACC^39^, which proceeds changes in the values of the targeted item^66^. Even though value-encoding areas, such as vmPFC and orbitofrontal cortices^68-70^, and cognitive-control-related areas, such as dlPFC^71,72^, might be engaged in later stages^48,73^, their role might not be as essential in initial nonreinforced learning^39,60^.

If PCC integrates behavioral tendencies, one should expect that PCC would be involved in guiding attention and initiating behavior regardless of external rewards and punishments. Remarkably, PCC, a central node in the default mode network, has been discussed as the neural hub that detects changes in the environment and motivates shifts in behavior^47^. Further, Leech and Sharp ^50^ argued that PCC plays a vital role in directing the focus of attention and supports internally-directed cognition.

Our VBM results show that PCC grey matter density predicts PCC BOLD activity changes. This finding may explain individual differences in responsiveness to behavioral modification training. In other words, participants who rely more heavily on behavioral tendencies in their daily life potentially have higher PCC grey matter density and are also more susceptible to AAT. This suggestion aligns with the idea that addiction might alter the importance and organization of neural activity in different brain areas^74^. However, this speculation should be treated cautiously before future studies replicate these results.

## Conclusion

In the current study, we challenged the assumption that value modification is necessary for altering choice behavior by investigating the effectiveness of AAT in facilitating choices of low-calorie food images. In line with other studies^13,14^, our results showed that AAT can effectively alter participants’ choice behavior. The observed behavioral modulations were related to alterations in indifference points and not subjective ratings, suggesting behavioral tendencies affected choice behavior while bypassing existing values. Our fMRI data revealed that the acceptance rates of low-calorie food items were correlated with PCC activity, which suggests PCC as the neural hub that integrates behavioral tendencies in decision-making processes. Finally, a correlation between BOLD activity changes and PCC grey matter density suggested a possible anatomical predisposition to behavioral modifications. Significantly, the current study indicates the possibility of affecting choice behavior regardless of value modification. This finding calls for a revision of existing frameworks used for understanding nonreinforced learning and value-based decision-making. Finally, our results offer a viable approach for improving health-related choices regardless of associated values, which is of great advantage for optimizing health-related decisions, such as food choices^20-23^.

## Method

### Participants and Stimuli

34 normal-weight (i.e., 18 < BMI < 25), right handed subjects (mean age = 25.14 years, SD = 4.01 years; mean BMI = 21.46, SD = 1.77) participated in the study. The sample size was chosen based on a priori power analysis with *α* = 0.05, 1 − *β* = 0.95 ^75,76^, and expected effect sizes of *Cohen’s f* = 0.35 (equivalent to 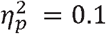 derived from previous studies^12-14,58^. The power analysis indicated that the total sample size *N* ≥ 29 were required. Participants did not report any history of mental illnesses. It has been reported that there are gender differences in metabolism^77^, eating behavior^78-80^, and neural responses to food cues^78^. Therefore, only female subjects were recruited for the study. Food intolerances or allergies and any diet restriction (i.e., being vegan or vegetarian) were exclusion criteria during recruitment. Prior to the experiment, all subjects were informed about the procedure and personal data handling, and their written consent was collected according to the declaration of Helsinki. Participation was compensated with 8€ per hour. The study was approved by the ethics committee of the University of Lübeck. One subject was excluded from data analyses completely, as she did not show any variance in the data (i.e., no accepted response in the food-choice task). Two more participants were excluded from analyses that required food ratings as they had more than 50% missed responses in the subjective rating task.

We selected 80 food images from the database of Blechert et al. ^53^, containing 40 low-(mean = 70.15kcal/100g, SD = 44.24 kcal/100g) and 40 high-calorie food items (m = 236.59 kcal/100g, SD = 59.24kcal/100g). Since the perception of sugar can interfere with and mask the perception of fat^81,82^, only savory items were selected. The selected food stimuli were divided into two stimulus stets, each containing 20 high- and 20 low-calorie items. To increase the training effect’s generalizability, one stimulus set was used exclusively for the pre- and post-tests and the other for AAT (i.e., the training session), making the results independent of the specific food items. The assignment of stimulus sets to tasks (i.e., test vs. training) was counterbalanced across participants. The use of different stimulus sets for training (i.e., AAT) and test (i.e., the choice and subjective rating tasks) ensured that the observed training effects are generalized over the stimuli, task, and context (i.e., in vs. outside of the MRI scanner).

### Choice and Subjective Rating Tasks

All tasks were programmed in MATLAB (r2020b; MathWorks Company) via Psychtoolbox v3^83^. In the subjective rating task, participants were asked to indicate how appealing they find a presented item on a continuous scale of 1-7, representing “not appealing at all” and “very appealing”, respectively, using a mouse. Each trial started with a fixation cross, randomly presented between 500-1000 ms. Afterward, a food item accompanied by the continuous Likert scale below the image was presented. Each trial would terminate either after the participants’ response or the maximum of 5000 ms. No response or responses after 5000 ms were considered as missed trials. Further, the order of images was completely randomized. Images (660×660 pixels) were presented against a black background.

In the food-choice task, participants indicated whether they were willing to consume a presented item or receive the equivalent amount of money instead. Two options (i.e., Yes and No) could be chosen by moving an MRI-compatible joystick (Fiber Optic Joystick, Current Designs) to the left or right. In order to avoid motor confounds on imaging data, the position (i.e., being on the left or right) of the “Yes” and “No” responses were counterbalanced among trials. Each trial was started by a fixation cross, randomly presented for 2, 4, 6, or 8 seconds (mean intertrial intervals = 4 s) to ensure hyperbolically distributed intertrial intervals^84^. Afterward, each stimulus was presented for 2000 ms, during which participants could respond. No response or responses after 2000 ms were considered as missed trials. Each image in the test set was presented four times during the food-choice task, yielding 160 trials in total. The task was split into four blocks of 40 trials, each consisting of 20 low- and 20 high-calorie images. The order of images was pseudorandomized so that each image would be presented only once during each block. The inter-block interval was 20 seconds. Images (660×660 pixels) were presented in a white frame against a black background.

### Approach-Avoidance Training

For AAT, participants were instructed to pull [towards their body] or push [away from their body] in response to the color of stimulus frames (i.e., blue or yellow) using an MRI-compatible joystick. Frame color to response assignment was counterbalanced across participants. For optimal approach and avoidance resemblance^54^, the employed AAT had an embedded zooming feature: pulling and pushing the joystick, depending on the approach or avoidance assignment, either increased or decreased the size of the presented image. In AAT, all low-and high-calorie stimuli were cued to be approached and avoided, respectively. Each picture in the training set was presented five times, yielding 200 trials in total. Further, AAT was split into five blocks of 40 trials, consisting of 20 low- and 20 high-calorie images. The order of images was pseudorandomized so that each image would be presented only once during each block. The trial structure (i.e., intertrial, inter-block intervals, and the response time) was similar to the food-choice task.

### Procedure

The study consisted of three sessions on three consecutive days. Prior to the study, participants were instructed to fast (i.e., do not eat and drink anything rather than water) and not consume alcohol or caffeine for at least 12 hours before each session. Every session started between 08:00 and 9:00 in the lab. In the beginning, participants received written instructions about the experimental procedure and the tasks of the respective day. On Days 1 and 3 (i.e., the pre and post-training sessions), the food-choice task was administered in the MRI scanner, followed by the subjective rating task, which was completed outside the scanner. Furthermore, on Day 2 (i.e., the training session), AAT was conducted outside the MRI scanner. Pre and post-training sessions lasted approximately one hour, and the AAT session took place in 30 minutes. At the end of each session, participants were offered a breakfast, where they could choose as many food items as they wanted from multiple available low- and high-calorie options. The number of the low- and high-calorie items that participants selected in pre- and post-session was recorded. All presented results are related to the comparisons of pre-vs. post-sessions.

### fMRI Acquisition and Preprocessing

Functional and anatomical images were acquired using a 3T Trio (Siemens) scanner equipped with a 12-channel head coil. In each of the four functional scanning runs, 127 T2*-weighted echo-planar images containing 33 slices, with 3 mm thickness and separated by a gap of 0.75 mm, were acquired. The order of acquisition was descending. Imaging parameters, resulting in an isotropic voxel size of 3 mm, were as follows: repetition time (TR), 2000 ms; echo time (TE), 30 ms; flip angle, 78°; matrix size, 64 × 64; field of view (FOV), 192 × 192 mm^2^. Further, a high-resolution T1-weighted magnetization prepared rapid gradient-echo image (MPRAGE) was collected for each subject. The parameters were as follows: TR, 1900 ms; TE, 2.52 ms; matrix size, 256 × 256; FOV, 256 × 256 mm^2^; 192 slices (1 mm thick); flip angle, 9°. Preprocessing, first-level, and group-level analysis of the functional data were conducted using SPM12 (The Wellcome Department of Imaging Neuroscience, Institute of Neurology, London, UK). During preprocessing, scans were spatially realigned, slice-time corrected, co-registered to their structural images, and subsequently normalized to the standard Montreal Neurological Institute (MNI) EPI template using deformation fields. Finally, images were smoothed using an 8-mm full-width at half-maximum (FWHM) Gaussian kernel. None of the participants moved more than 3 mm/rad within each run.

### Statistical Analysis

All behavioral statistical analyses were conducted via R programming language (http://www.R-project.org/). Only correct responses were used for statistical analyses. The results of the food-choice and subjective rating tasks were analyzed using a 2 × 2 repeated measure ANOVA, with Session (pre vs. post) and Calorie (high vs. low) as within-subject factors. Further, indifference points were calculated using logistic regression modeling. Indifference points in binary choices are referred to estimated positions where agents might accept or reject an item with similar probability^51,52^. For each participant in each condition, choices were entered into the model as a binary input (i.e., yes = 1, no = 0) and subjective ratings as a continuous predictor. The model’s output represents the probability of choosing an item giving the subjective rating for that item, as described in equation 1.

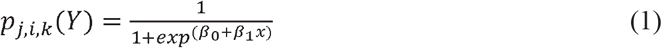

In equation 1, *x* designates subjective rating, *Y* choice, *j* participant number, *i* session (e.g., pre-training), *k* calorie content (e.g., low-calorie), and finally *β*_0_ and *β*_1_ are the parameters of the model. For each of the participants at each condition, the indifference points were defined as the subjective rating that predicts choosing an item with the probability of 50%. For five participants, the rating associated with 50% was outside acceptable boundaries (i.e., [1,7]); therefore, these participants were eliminated in the final analysis of indifference points. The calculated indifference points were entered in a 2 × 2 repeated measure ANOVA similar to the one used for the subjective rating and food-choice tasks. All the behavioral data were checked to conform to ANOVA’s assumptions: normal distributions, no extreme outliers [i.e., outside *Quartile* 1(*Q*1) − 3 * *interquartile*(*IQR*), *Q*3 + 3 * *IQR*], and linearity of relations.

For participants’ breakfast choices, the output (i.e., the count of selected low- and high-calorie food items) had a quasi-Poisson instead of a Gaussian distribution. Therefore, for analyzing the breakfast data, we utilized quasi-Poisson mixed models with Participant included as a random and Session (pre vs. post) and Calorie (high vs. low) as fixed effects. For calculating and analyzing the model, the MASS package in R was used^85^. The statistical inference was made based on chi-square tests.

The statistical analyses of fMRI data were conducted using MATLAB (r2020b; MathWorks Company), SPM12 (The Wellcome Department of Imaging Neuroscience, Institute of Neurology, London, UK), and R programming language (http://www.R-project.org/). The effects of interest were calculated for each participant and session using general linear models (GLM), including all four runs. In GLM1, the food-choice task trials were assigned to four event-related conditions: accepted and rejected items of low (n = 80) and high-calorie images (n = 80). The resulted vectors were convolved with a canonical hemodynamic response function. Further, we used a high-pass filter with a 180 Hz cutoff and an explicit brain mask. For finding functional ROIs, *p* values were corrected for multiple comparisons using the family-wise error correction at the cluster level (*FWE*_*c*_). Based on the suggestions of Woo et al. ^86^, we chose a high primary threshold (i.e., *p* < 0.001, *contigueous voxels* > 40) to enhance spatial localization and interpretability. This approach has been discussed to provide the best balance between the type I and II errors in fMRI studies^56,86^. Extracted neural activities of selected ROIs were assessed using a 2 × 2 × 2 repeated measure ANOVA, with Session (pre vs. post), Calorie (high vs. low), and Decision (accepted vs. reject) as within-subject factors. For calculating correlations between neural activities and behavioral measures, the repeated-measure correlation package (rmcorr) in R was used^87^.

For assessing the parametric modulation of blood oxygenation level-dependent (BOLD) responses by subjective ratings, GML2 was devised in which ratings of presented food items were included in the model. To ensure that multicollinearity will not affect our results, the subjective ratings were mean-centered for each participant and condition (i.e., low and high-calorie food items)^88^. In GLM2 pre and post-training sessions were combined, and therefore, the design matrix was reduced to 2 × 2: two levels of Calorie and two levels of Decision. The rest of the model was identical to GLM1.

Further, to assess whether there can be a predisposition to AAT, we conducted voxel-based morphometry (VBM) using the computational anatomy toolbox (CAT12). For VBM, T1 images are spatially normalized using geodesic shooting templates^89^ and segmented into gray matter (GM), white matter (WM), and cerebrospinal fluid (CSF). Total intracranial volumes (TIV) were calculated and used as a nuisance regressor in VBM-GLM. Afterward, images were smoothed using an 8-mm FWHM. Finally, the extracted grey matter densities of the selected regions of interest (ROI) were correlated with the BOLD activity of those same regions. Since we did not assume a linear relationship between BOLD activity and grey matter density, instead of Pearson’s correlation, ranked Spearman’s correlation was assessed.

## Acknowledgment

We thank Riccardo Barbieri for his immense help during data collection of the experiment. SQP was supported by the German Federal Ministry of Education and Research and the Federal State of Brandenburg (BMBF), DZD grant 82DZD03D03- and DZD Grant 82DZD03C2G and by German Research Foundation Grant/Award Number INST 392/125 (SFB TR134 C7) and PA2682/1-1. KG was supported by the German Research Foundation (DFG, EXC 2002/1 “Science of Intelligence” – project number 390523135).

## References

1 Romieu, I. et al. Energy balance and obesity: what are the main drivers? Cancer Causes Control 28, 247–258, doi:10.1007/s10552-017-0869-z. (2017).

2 Forouzanfar, M. H. et al. Global, regional, and national comparative risk assessment of 79 behavioural, environmental and occupational, and metabolic risks or clusters of risks in 188 countries, 1990–2013: a systematic analysis for the Global Burden of Disease Study 2013. The Lancet 386, 2287–2323, doi:10.1016/s0140-6736(15)00128-2 (2015).

3 Ebbeling, C. B. et al. Compensation for energy intake from fast food among overweight and lean adolescents. JAMA 291, 2828–2833 (2004).

4 Cureau, F. V., Sparrenberger, K., Bloch, K. V., Ekelund, U. & Schaan, B. D. Associations of multiple unhealthy lifestyle behaviors with overweight/obesity and abdominal obesity among Brazilian adolescents: A country-wide survey. Nutr. Metab. Cardiovasc. Dis. 28, 765–774, doi:10.1016/j.numecd.2018.04.012 (2018).

5 Clark, M. A., Springmann, M., Hill, J. & Tilman, D. Multiple health and environmental impacts of foods. Proc. Natl. Acad. Sci. U. S. A., doi:10.1073/pnas.1906908116 (2019).

6 Mason, P. & Lang, T. Sustainable diets: how ecological nutrition can transform consumption and the food system. (Routledge, 2015).

7 Willett, W. et al. Food in the Anthropocene: the EAT–Lancet Commission on healthy diets from sustainable food systems. The Lancet 393, 447–492, doi:10.1016/s0140-6736(18)31788-4 (2019).

8 Balleine, B., Daw, N. & O’Doherty, J. (New York: Academic Press.[Google Scholar], 2008).

9 Balleine, B. W. & Dickinson, A. Goal-directed instrumental action: contingency and incentive learning and their cortical substrates. Neuropharmacology 37, 407–419, doi:10.1016/s0028-3908(98)00033-1 (1998).

10 Balleine, B. W. & O’Doherty, J. P. Human and rodent homologies in action control: corticostriatal determinants of goal-directed and habitual action. Neuropsychopharmacology 35, 48–69, doi:10.1038/npp.2009.131 (2010).

11 Rangel, A., Camerer, C. & Montague, P. R. A framework for studying the neurobiology of value-based decision making. Nat. Rev. Neurosci. 9, 545–556, doi:10.1038/nrn2357 (2008).

12 Schumacher, S. E., Kemps, E. & Tiggemann, M. Bias modification training can alter approach bias and chocolate consumption. Appetite 96, 219–224, doi:10.1016/j.appet.2015.09.014 (2016).

13 Mehl, N., Mueller-Wieland, L., Mathar, D. & Horstmann, A. Retraining automatic action tendencies in obesity. Physiol. Behav. 192, 50–58, doi:10.1016/j.physbeh.2018.03.031 (2018).

14 Kakoschke, N., Kemps, E. & Tiggemann, M. The effect of combined avoidance and control training on implicit food evaluation and choice. J. Behav. Ther. Exp. Psychiatry 55, 99–105, doi:10.1016/j.jbtep.2017.01.002 (2017).

15 Becker, D., Jostmann, N. B., Wiers, R. W. & Holland, R. W. Approach avoidance training in the eating domain: testing the effectiveness across three single session studies. Appetite 85, 58–65, doi:10.1016/j.appet.2014.11.017 (2015).

16 Bakkour, A. et al. Mechanisms of Choice Behavior Shift Using Cue-approach Training. Front. Psychol. 7, 421, doi:10.3389/fpsyg.2016.00421 (2016).

17 Schonberg, T. et al. Changing value through cued approach: an automatic mechanism of behavior change. Nat. Neurosci. 17, 625–630, doi:10.1038/nn.3673 (2014).

18 Schonberg, T. & Katz, L. N. A Neural Pathway for Nonreinforced Preference Change. Trends Cogn. Sci. 24, 504–514, doi:10.1016/j.tics.2020.04.002 (2020).

19 Knudsen, E. B. & Wallis, J. D. Taking stock of value in the orbitofrontal cortex. Nat. Rev. Neurosci., doi:10.1038/s41583-022-00589-2 (2022).

20 Bongers, P. et al. Being impulsive and obese increases susceptibility to speeded detection of high-calorie foods. Health Psychol. 34, 677–685, doi:10.1037/hea0000167 (2015).

21 Guerrieri, R., Nederkoorn, C. & Jansen, A. The interaction between impulsivity and a varied food environment: its influence on food intake and overweight. Int. J. Obes. (Lond.) 32, 708–714, doi:10.1038/sj.ijo.0803770 (2008).

22 Nederkoorn, C., Guerrieri, R., Havermans, R. C., Roefs, A. & Jansen, A. The interactive effect of hunger and impulsivity on food intake and purchase in a virtual supermarket. Int. J. Obes. (Lond.) 33, 905–912, doi:10.1038/ijo.2009.98 (2009).

23 Nederkoorn, C., Houben, K., Hofmann, W., Roefs, A. & Jansen, A. Control yourself or just eat what you like? Weight gain over a year is predicted by an interactive effect of response inhibition and implicit preference for snack foods. Health Psychol. 29, 389–393, doi:10.1037/a0019921 (2010).

24 Thrailkill, E. A., Michaud, N. L. & Bouton, M. E. Reinforcer predictability and stimulus salience promote discriminated habit learning. J Exp Psychol Anim Learn Cogn 47, 183–199, doi:10.1037/xan0000285 (2021).

25 Colwill, R. M. & Delamater, B. A. An associative analysis of instrumental biconditional discrimination learning. Anim. Learn. Behav. 23, 218–233, doi:10.3758/bf03199937 (1995).

26 Colwill, R. M. An Associative Analysis of Instrumental Learning. Current Directions in Psychological Science 2, 111–116, doi:10.1111/1467-8721.ep10772598 (1993).

27 Bradfield, L. A. & Balleine, B. W. Hierarchical and binary associations compete for behavioral control during instrumental biconditional discrimination. J. Exp. Psychol. Anim. Behav. Process. 39, 2–13, doi:10.1037/a0030941 (2013).

28 Hershberger, W. A. An approach through the looking-glass. Anim. Learn. Behav. 14, 443–451, doi:10.3758/bf03200092 (1986).

29 Strack, F. & Deutsch, R. Reflective and impulsive determinants of social behavior. Pers. Soc. Psychol. Rev. 8, 220–247, doi:10.1207/s15327957pspr0803_1 (2004).

30 Cooper, R. & Shallice, T. Contention scheduling and the control of routine activities. Cogn. Neuropsychol. 17, 297–338, doi:10.1080/026432900380427 (2000).

31 Norman, D. A. & Shallice, T. in Consciousness and Self-Regulation Ch. Chapter 1, 1-18 (1986).

32 Diamond, A. Executive functions. Annu. Rev. Psychol. 64, 135–168, doi:10.1146/annurev-psych-113011-143750 (2013).

33 Botvinick, M. & Braver, T. Motivation and cognitive control: from behavior to neural mechanism. Annu. Rev. Psychol. 66, 83–113, doi:10.1146/annurev-psych-010814-015044 (2015).

34 Shenhav, A. et al. Toward a Rational and Mechanistic Account of Mental Effort. Annu. Rev. Neurosci. 40, 99–124, doi:10.1146/annurev-neuro-072116-031526 (2017).

35 Froehlich, E. et al. A short humorous intervention protects against subsequent psychological stress and attenuates cortisol levels without affecting attention. Sci. Rep. 11, 7284, doi:10.1038/s41598-021-86527-1 (2021).

36 Zahedi, A., Luczak, A. & Sommer, W. Modification of food preferences by posthypnotic suggestions: An event-related brain potential study. Appetite 151, 104713, doi:10.1016/j.appet.2020.104713 (2020).

37 Wiers, R. W., Eberl, C., Rinck, M., Becker, E. S. & Lindenmeyer, J. Retraining automatic action tendencies changes alcoholic patients’ approach bias for alcohol and improves treatment outcome. Psychol. Sci. 22, 490–497, doi:10.1177/0956797611400615 (2011).

38 Kakoschke, N., Kemps, E. & Tiggemann, M. Approach bias modification training and consumption: A review of the literature. Addict. Behav. 64, 21–28, doi:10.1016/j.addbeh.2016.08.007 (2017).

39 Mehl, N., Morys, F., Villringer, A. & Horstmann, A. Unhealthy yet Avoidable-How Cognitive Bias Modification Alters Behavioral and Brain Responses to Food Cues in Individuals with Obesity. Nutrients 11, doi:10.3390/nu11040874 (2019).

40 Wiers, C. E. et al. Effects of cognitive bias modification training on neural signatures of alcohol approach tendencies in male alcohol-dependent patients. Addict. Biol. 20, 990–999, doi:10.1111/adb.12221 (2015).

41 Bartra, O., McGuire, J. T. & Kable, J. W. The valuation system: a coordinate-based meta-analysis of BOLD fMRI experiments examining neural correlates of subjective value. NeuroImage 76, 412–427, doi:10.1016/j.neuroimage.2013.02.063 (2013).

42 Kable, J. W. & Glimcher, P. W. The Neurobiology of Decision: Consensus and Controversy. Neuron 63, 733–745, doi:10.1016/j.neuron.2009.09.003 (2009).

43 Kable, J. W. & Glimcher, P. W. The neural correlates of subjective value during intertemporal choice. Nat. Neurosci. 10, 1625–1633, doi:10.1038/nn2007 (2007).

44 Paulus, M. P. & Frank, L. R. Ventromedial prefrontal cortex activation is critical for preference judgments. Neuroreport 14, 1311–1315, doi:10.1097/01.wnr.0000078543.07662.02 (2003).

45 Plassmann, H., O’Doherty, J. & Rangel, A. Orbitofrontal cortex encodes willingness to pay in everyday economic transactions. J. Neurosci. 27, 9984–9988, doi:10.1523/JNEUROSCI.2131-07.2007 (2007).

46 Plassmann, H., O’Doherty, J. P. & Rangel, A. Appetitive and aversive goal values are encoded in the medial orbitofrontal cortex at the time of decision making. J. Neurosci. 30, 10799–10808, doi:10.1523/JNEUROSCI.0788-10.2010 (2010).

47 Pearson, J. M., Heilbronner, S. R., Barack, D. L., Hayden, B. Y. & Platt, M. L. Posterior cingulate cortex: adapting behavior to a changing world. Trends Cogn. Sci. 15, 143–151, doi:10.1016/j.tics.2011.02.002 (2011).

48 Cousijn, J. et al. Approach-bias predicts development of cannabis problem severity in heavy cannabis users: results from a prospective FMRI study. PLoS One 7, e42394, doi:10.1371/journal.pone.0042394 (2012).

49 Ko, C. H. et al. Brain correlates of craving for online gaming under cue exposure in subjects with Internet gaming addiction and in remitted subjects. Addict. Biol. 18, 559–569, doi:10.1111/j.1369-1600.2011.00405.x (2013).

50 Leech, R. & Sharp, D. J. The role of the posterior cingulate cortex in cognition and disease. Brain 137, 12–32, doi:10.1093/brain/awt162 (2014).

51 McKerchar, T. L. et al. A comparison of four models of delay discounting in humans. Behav. Processes 81, 256–259, doi:10.1016/j.beproc.2008.12.017 (2009).

52 Scherbaum, S., Dshemuchadse, M. & Goschke, T. Building a bridge into the future: dynamic connectionist modeling as an integrative tool for research on intertemporal choice. Front. Psychol. 3, 514, doi:10.3389/fpsyg.2012.00514 (2012).

53 Blechert, J., Meule, A., Busch, N. A. & Ohla, K. Food-pics: an image database for experimental research on eating and appetite. Front. Psychol. 5, 617, doi:10.3389/fpsyg.2014.00617 (2014).

54 Rinck, M. & Becker, E. S. Approach and avoidance in fear of spiders. J. Behav. Ther. Exp. Psychiatry 38, 105–120, doi:10.1016/j.jbtep.2006.10.001 (2007).

55 Meule, A. & Kubler, A. Double trouble. Trait food craving and impulsivity interactively predict food-cue affected behavioral inhibition. Appetite 79, 174–182, doi:10.1016/j.appet.2014.04.014 (2014).

56 Lieberman, M. D. & Cunningham, W. A. Type I and Type II error concerns in fMRI research: re-balancing the scale. Soc. Cogn. Affect. Neurosci. 4, 423–428, doi:10.1093/scan/nsp052 (2009).

57 Dickson, H., Kavanagh, D. J. & MacLeod, C. The pulling power of chocolate: Effects of approach-avoidance training on approach bias and consumption. Appetite 99, 46–51, doi:10.1016/j.appet.2015.12.026 (2016).

58 Jones, A., Hardman, C. A., Lawrence, N. & Field, M. Cognitive training as a potential treatment for overweight and obesity: A critical review of the evidence. Appetite 124, 50–67, doi:10.1016/j.appet.2017.05.032 (2018).

59 Bakkour, A., Lewis-Peacock, J. A., Poldrack, R. A. & Schonberg, T. Neural mechanisms of cue-approach training. NeuroImage 151, 92–104, doi:10.1016/j.neuroimage.2016.09.059 (2017).

60 Aridan, N., Pelletier, G., Fellows, L. K. & Schonberg, T. Is ventromedial prefrontal cortex critical for behavior change without external reinforcement? Neuropsychologia 124, 208–215, doi:10.1016/j.neuropsychologia.2018.12.008 (2019).

61 Botvinik-Nezer, R., Salomon, T. & Schonberg, T. Enhanced Bottom-Up and Reduced Top-Down fMRI Activity Is Related to Long-Lasting Nonreinforced Behavioral Change. Cereb. Cortex 30, 858–874, doi:10.1093/cercor/bhz132 (2020).

62 Levey, A. B. & Martin, I. Classical conditioning of human ‘evaluative’ responses. Behaviour Research and Therapy 13, 221–226, doi:10.1016/0005-7967(75)90026-1 (1975).

63 Hutter, M. & Rothermund, K. Automatic processes in evaluative learning. Cogn. Emot. 34, 1–20, doi:10.1080/02699931.2019.1709315 (2020).

64 Hofmann, W., De Houwer, J., Perugini, M., Baeyens, F. & Crombez, G. Evaluative conditioning in humans: a meta-analysis. Psychol. Bull. 136, 390–421, doi:10.1037/a0018916 (2010).

65 Corneille, O. & Stahl, C. Associative Attitude Learning: A Closer Look at Evidence and How It Relates to Attitude Models. Pers. Soc. Psychol. Rev. 23, 161–189, doi:10.1177/1088868318763261 (2019).

66 Voigt, K., Murawski, C., Speer, S. & Bode, S. Hard Decisions Shape the Neural Coding of Preferences. J. Neurosci. 39, 718–726, doi:10.1523/JNEUROSCI.1681-18.2018 (2019).

67 Zhou, Y. & Freedman, D. J. Posterior parietal cortex plays a causal role in perceptual and categorical decisions. Science 365, 180–185, doi:10.1126/science.aaw8347 (2019).

68 Baumgartner, T., Knoch, D., Hotz, P., Eisenegger, C. & Fehr, E. Dorsolateral and ventromedial prefrontal cortex orchestrate normative choice. Nat. Neurosci. 14, 1468–1474, doi:10.1038/nn.2933 (2011).

69 Smith, D. V., Clithero, J. A., Boltuck, S. E. & Huettel, S. A. Functional connectivity with ventromedial prefrontal cortex reflects subjective value for social rewards. Soc. Cogn. Affect. Neurosci. 9, 2017–2025, doi:10.1093/scan/nsu005 (2014).

70 Smith, D. V. et al. Distinct value signals in anterior and posterior ventromedial prefrontal cortex. J. Neurosci. 30, 2490–2495, doi:10.1523/JNEUROSCI.3319-09.2010 (2010).

71 Yan, Y., Wei, R., Zhang, Q., Jin, Z. & Li, L. Differential roles of the dorsal prefrontal and posterior parietal cortices in visual search: a TMS study. Sci. Rep. 6, 30300, doi:10.1038/srep30300 (2016).

72 Wang, M. et al. Evaluating the causal contribution of fronto-parietal cortices to the control of the bottom-up and top-down visual attention using fMRI-guided TMS. Cortex 126, 200–212, doi:10.1016/j.cortex.2020.01.005 (2020).

73 Wiers, C. E. et al. Neural Correlates of Alcohol-Approach Bias in Alcohol Addiction: the Spirit is Willing but the Flesh is Weak for Spirits. Neuropsychopharmacology 39, 688–697, doi:10.1038/npp.2013.252 (2013).

74 Volkow, N. D. et al. Addiction: decreased reward sensitivity and increased expectation sensitivity conspire to overwhelm the brain’s control circuit. Bioessays 32, 748–755, doi:10.1002/bies.201000042 (2010).

75 Cohen, J. Statistical power analysis for the behavioral sciences {Ind cd. Hillsdale. NJ: Erihaum (1988).

76 Cohen, J. Statistical Power Analysis. Current Directions in Psychological Science 1, 98–101, doi:10.1111/1467-8721.ep10768783 (2016).

77 Dionne, I., Despres, J. P., Bouchard, C. & Tremblay, A. Gender difference in the effect of body composition on energy metabolism. Int. J. Obes. Relat. Metab. Disord. 23, 312–319, doi:10.1038/sj.ijo.0800820 (1999).

78 Manippa, V., Padulo, C., van der Laan, L. N. & Brancucci, A. Gender Differences in Food Choice: Effects of Superior Temporal Sulcus Stimulation. Front. Hum. Neurosci. 11, 597, doi:10.3389/fnhum.2017.00597 (2017).

79 Rolls, B. J., Fedoroff, I. C. & Guthrie, J. F. Gender differences in eating behavior and body weight regulation. Health Psychol. 10, 133–142, doi:10.1037/0278-6133.10.2.133 (1991).

80 Wardle, J. et al. Gender differences in food choice: the contribution of health beliefs and dieting. Ann. Behav. Med. 27, 107–116, doi:10.1207/s15324796abm2702_5 (2004).

81 Bolhuis, D. P., Costanzo, A. & Keast, R. S. J. Preference and perception of fat in salty and sweet foods. Food Qual. Prefer. 64, 131–137, doi:10.1016/j.foodqual.2017.09.016 (2018).

82 Drewnowski, A. & Schwartz, M. Invisible fats: Sensory assessment of sugar/fat mixtures. Appetite 14, 203–217, doi:10.1016/0195-6663(90)90088-p (1990).

83 Brainard, D. H. The Psychophysics Toolbox. Spat. Vis. 10, 433–436, doi:10.1163/156856897x00357 (1997).

84 Miezin, F. M., Maccotta, L., Ollinger, J. M., Petersen, S. E. & Buckner, R. L. Characterizing the hemodynamic response: effects of presentation rate, sampling procedure, and the possibility of ordering brain activity based on relative timing. NeuroImage 11, 735–759, doi:10.1006/nimg.2000.0568 (2000).

85 Ripley, B. et al. Package ‘mass’. Cran r 538, 113–120 (2013).

86 Woo, C. W., Krishnan, A. & Wager, T. D. Cluster-extent based thresholding in fMRI analyses: pitfalls and recommendations. NeuroImage 91, 412–419, doi:10.1016/j.neuroimage.2013.12.058 (2014).

87 Bakdash, J. Z. & Marusich, L. R. Repeated Measures Correlation. Front. Psychol. 8, 456, doi:10.3389/fpsyg.2017.00456 (2017).

88 Iacobucci, D., Schneider, M. J., Popovich, D. L. & Bakamitsos, G. A. Mean centering helps alleviate “micro” but not “macro” multicollinearity. Behav. Res. Methods 48, 1308–1317, doi:10.3758/s13428-015-0624-x (2016).

89 Ashburner, J. & Friston, K. J. Diffeomorphic registration using geodesic shooting and Gauss-Newton optimisation. NeuroImage 55, 954–967, doi:10.1016/j.neuroimage.2010.12.049 (2011).

